# Fitting and comparison of calcium-calmodulin kinetic schemes to a common data set using non-linear mixed effects modelling

**DOI:** 10.1101/2024.10.28.620577

**Authors:** Domas Linkevicius, Angus Chadwick, Guido C. Faas, Melanie I. Stefan, David C. Sterratt

## Abstract

Calmodulin is a calcium binding protein that is essential in calcium signalling in the brain. There are many computational models of calcium-calmodulin binding that capture various calmodulin features. However, existing models have generally been fit to different data sets, with some publications not reporting their training and validation performance. Moreover, there is no model comparison using a common benchmark data set as is common practice in other modeling domains. Finally, some calmodulin models have been fit as a part of a larger kinetic scheme, which may have resulted in parameters being underdetermined.

We address these three limitations of previous models by fitting the published calcium-calmodulin schemes to a common calcium-calmodulin data set comprising of equlibrium data from Shifman et al. and dynamical data from Faas et al. Due to technical limitations, the amount of uncaged calcium in Faas et al. data could not be predicted with certainty. To find good parameter fits, despite this uncertainty, we used non-linear mixed effects modelling as implemented in the `Pumas.jl` package.

The Akaike information criterion values for our reaction rates were significantly lower than for the published parameters, indicating that the published parameters are suboptimal. Moreover, there were significant differences in calmodulin activation, both between the schemes and between our reaction rate and those previously published. A kinetic scheme with independent lobes and unique, rather than identical, binding sites fit the data best. Our results support two hypotheses: (1) partially bound calmodulin is important in cellular signalling; (2) calcium binding sites within a calmodulin lobe are kinetically distinct rather than identical. We conclude that more attention should be given to validation and comparison of models of individual molecules.

**Author summary:** Learning and memory depend on changes in synapses, the connections between nerve cells. To understand how learning and memory work, it is important to understand how the proteins involved in these changes are regulated. Computational modelling of biochemical reactions has been used to understand the regulation processes involved in learning and memory. However, computational models often rely on simplifications of biochemical reaction schemes, and it can be difficult to tell which simplifications are “the best”, i.e. capture important aspects of a protein’s behaviour. It is also difficult to estimate the best model parameters, such as binding reaction rates or binding strengths. Modellers often base their estimates on known experimental results, but often not in a structured way.

We examine and compare a variety of computational models of calmodulin, a protein necessary for changes in synaptic strength. We apply a new approach to infer the model parameters that best fit two published datasets, and compare our parameters to previously published ones. We find that some simplified reaction schemes are more successful than others in capturing the activation pattern of calmodulin. We also find that the parameter sets found through our approach outperform previously published parameter sets in fitting the experimental data.

## Introduction

Calmodulin is among the most important calcium binding proteins in the brain. It is essential in the translation of intracellular Ca^2+^ signals to downstream processes, such as gene regulation, protein activation, metabolic regulation and synaptic plasticity [1, 2]. Calcium signals can provide information both via their amplitude (nanomolar to millimolar) and via their duration (microseconds to hours) [1, 2]. Calcium-calmodulin binding kinetics underlie the translation of Ca^2+^ signals, therefore correct kinetic models of binding are an important aspect in studying calcium signalling in the brain.

Calmodulin’s signalling properties arise from its structure – it comprises a 148 amino acid residue polypeptide with four EF hands divided into C and N lobes capable of binding two calcium ions per lobe [3]. It can adopt many conformational states, especially when bound to different molecules [4]. Moreover, calmodulin lobes have been reported to differ in their kinetics and affinity for Ca^2+^– the N lobe binding faster with lower affinity and the C lobe binding slower with higher affinity [5–7].

There are at least 19 published computational models of synapses that include various models of calmodulin in their chemical reaction network [8]. The published kinetic schemes describing calcium-calmodulin binding vary significantly in the number of calmodulin features they capture. For example, some calmodulin models do not have independent lobes [9–11] while others do [5, 12, 13]. Some schemes are event-based – only concerned about the Ca^2+^ binding events [10, 11], whereas others explicitly indicate which lobe and/or site is being bound to [5, 12, 13]. Moreover, some models assume that two Ca^2+^ ions bind to a lobe at the same time [9, 10, 12], others leave this dependent on reaction rate constants [5, 11, 13]. Some models assume that Ca^2+^ binding sites within a lobe are unique [11, 13] while others assume that they are non-unique [5, 12]. Finally, some models include details such as calmodulin conformational states [14].

Most current computational calmodulin models suffer from three limitations. First of all, different models have generally been tuned to different data sets, making their relative performance difficult to compare. Secondly, most models have not been cross-validated, making their generalization performance uncertain. Thirdly, some models have been tuned as a part of a larger scheme, e.g. including CaMKII, potentially making calcium-calmodulin binding parameters underdetermined. We discuss the sources of data to which calcium-calmodulin models we investigate have been fit in the Methods section, therefore we will next elaborate on the second and the third limitations.

The second limitation relates to cross-validation, a crucial step in the process of parameter inference used to establish model performance outside of the training data and to avoid overfitting (see page 241 in [15]). Ideally different models are cross-validated on a single data set across publications using consistent quantitative metrics. For example, the MNIST data set [16] is used to compare the error rate of image processing models. Given the lack of rich open access calmodulin data sets, none of the published calmodulin models were quantitatively cross-validated during development. At best, publications that contain calmodulin kinetic schemes include some indication (usually visual, rather than quantitative) of performance compared to the source of data they are being tuned to. However, this is not a rigorous way of ensuring that a model will perform well outside of the training data, leaving the generalization performance uncertain.

The third limitation is that some calmodulin schemes have been tuned to data from experiments that include other calmodulin-binding molecules, with large numbers of reaction rate constants that have to be fit, for example the calcium-calmodulin-CaMKII cascade [13]. Systems biology models naturally exhibit sloppiness [17], which tends to get more pronounced with an increasing number parameters being fit, resulting in loosely constrained parameter values. It is often possible to trade off between reaction rate constants: a calcium-calmodulin-CaMKII cascade being fit to CaMKII activity measurements may fit data better if Ca^2+^ binds to calmodulin with higher affinity or calcium-calmodulin binds to CaMKII with higher affinity, or some mix of the two. Moreover, a similar trade-off is possible between the binding sites and/or lobes even with only a calcium-calmodulin cascade. Because of these trade-offs, the true reaction rates might be completely different to the ones obtained via the fitting procedures. Some publications attempt to test for such parameter sloppiness via sensitivity analyses [18] or by calculating the eigenvalues of the Hessian [17], but it usually is too difficult to test the parameter combinations in a sufficiently dense and wide manner to ensure that the reaction rate constants are not under-determined.

Calmodulin models, in particular some of the simpler ones [9, 12], have been used to investigate calmodulin interactions with other molecules [18–20] and in complex chemical reaction networks [9, 21–23] to model higher order phenomena occurring in neurons, e.g. synaptic plasticity. However, given the aforementioned model limitations, it is important to scrutinize the previous modelling work, its basic assumptions, and to check whether the assumptions made in previous work hold when tested under more rigorous conditions, with powerful methods using richer data sets.

We address the three aforementioned limitations of existing calmodulin models by using a common data set where the only free kinetic parameters are calcium-calmodulin binding rates. The common data set comprises subsets of data from Faas et al. [5] and Shifman et al. [11]. Faas et al. [5] contains time-series of fluorescence measurements after laser-induced Ca^2+^ uncaging and therefore is informative about calmodulin dynamics. In contrast, Shifman et al. [11] contains measurements of calmodulin properties at equilibrium. To deal with incomplete experimental control of the amount of calcium uncaged by a laser flash in Faas et al. [5] we use the novel and highly efficient non-linear mixed effects model fitting algorithms implemented in `Pumas.jl` [24]. NLME is a hierarchical modeling framework that can deal with phenomena where there are constant intra-individual parameters, but significant inter-individual variability due to individual level parameters [24, 25].

We use the common data set to fit reaction rates from scratch and compare our results to the reaction rates in the literature. By calculating the Akaike information criterion (AIC) [26] values for both our and the published reaction rates we show that the published reaction rates are suboptimal. Moreover, using the same criterion, we show that some kinetic schemes are suboptimal and fail to fit calmodulin dynamics and equilibrium behaviour at the same time. We then compare the Ca^2+^ signal integration properties of different calmodulin schemes when either the published reaction rates or the ones determined by our approach are used. We show that there are significant differences in calmodulin calcium integration properties when using the suboptimal published reaction rates. Similarly, we show that the models using suboptimal calmodulin schemes display qualitatively different calcium integration behaviour compared to better performing schemes. Finally, we calculate the partial rank correlations between the reaction rate constants that we fit and show that for some calmodulin schemes our parameter fits are highly correlated which is indicative of parameter sloppiness or underdetermination.

Our results highlight that a sufficiently expressive calmodulin model structure is essential for capturing both calmodulin dynamics and equilibrium behaviour. Moreover, we conclude that, given the suboptimality of the previously published parameter sets, arguments and findings built on these models may warrant re-visiting.

## Methods

### Data

#### Dynamical calmodulin data

We use the calcium uncaging data from Faas et al. [5], in which different concentration mixes of the fluorescent Ca^2+^ indicator OGB-5N, the light sensitive Ca^2+^ chelator DM-nitrophen (DMn), calmodulin and titrated free Ca^2+^ were used to make seven different groups of solutions A–G (see S2 Table for specific concentrations). For different batches of each group of solutions, a sequence of laser pulses of increasing strength was used to induce Ca^2+^ uncaging from DMn while OGB-5N fluorescence was observed at 35°C. The stronger the pulse, the more calcium is released. Due to technical limitations, it was not possible to predict the amount of released calcium for each laser pulse precisely. We elaborate on how we model the fraction of uncaged Ca^2+^ below. Fig 1 shows the fluorescence time courses for three of the seven groups – A, B and G – and different uncaging laser strengths. We use a subset of the data and split it into training, validation and test data sets (see Supplemental Text S1 Appendix for more information).

**Fig 1.**
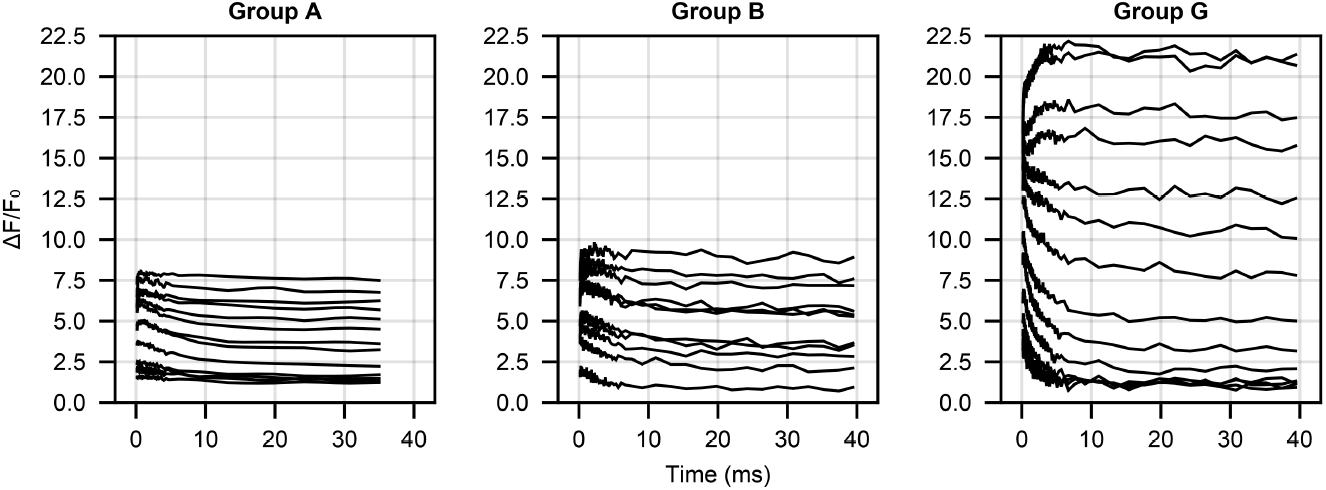
Relative fluorescence (∆*F/F*_0_) time-series data from [5] for three different initial condition groups (A, B and G). Different lines within a plot are due to different laser uncaging strength (the higher the laser strength, the larger the ∆*F/F*_0_ value).

#### Calmodulin equilibrium data

Steady-state calcium-calmodulin binding came from an experiment in Shifman et al. [11], which measured the number of Ca^2+^ ions bound per calmodulin molecule at different free Ca^2+^ concentrations (their Fig 1B). Their experimental chamber contained a fluorescent indicator Fluo4FF (5*μ*M), calmodulin (5*μ*M) and a varying amount of free Ca^2+^. The amount of free Ca^2+^ was titrated until a required concentration (between approximately 10^−7^M and 5.5 *×* 10^−5^M) was reached. We used a digital tool (https://automeris.io/WebPlotDigitizer/) to extract this data from their plots, giving the 107 points shown in Fig 2. This data was obtained at 25°C but calmodulin does not show significant temperature dependent changes in equilibrium behaviour [27], so we do not adjust for temperature dependent changes in calmodulin kinetics.

**Fig 2.**
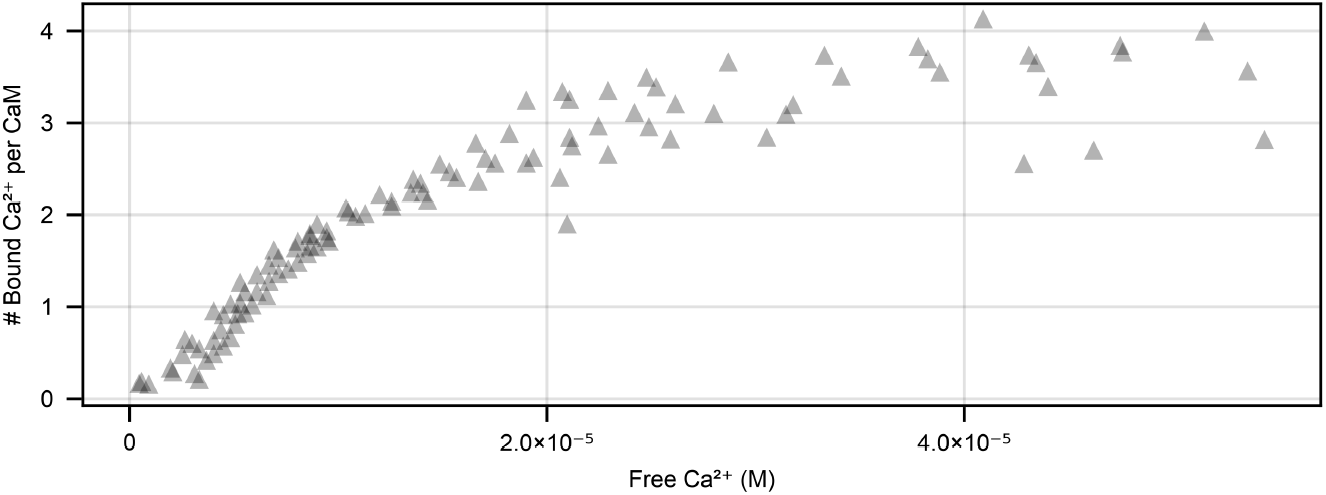
Equilibrium measurements of the number of Ca^2+^ ions per calmodulin molecule from [11]. Experiments were done using 5μM Fluo4FF and 5μM calmodulin.

### Kinetic schemes and published reaction rates

We investigated six different calcium-calmodulin binding schemes from the literature that span the complexity of the most commonly used calmodulin models (Fig 3). The reference we give for a scheme may be its original source, or a source that is frequently cited for the scheme. There are more complex published calmodulin schemes that we did not use [14], because they would be prohibitively computationally expensive to fit.

**Fig 3.**
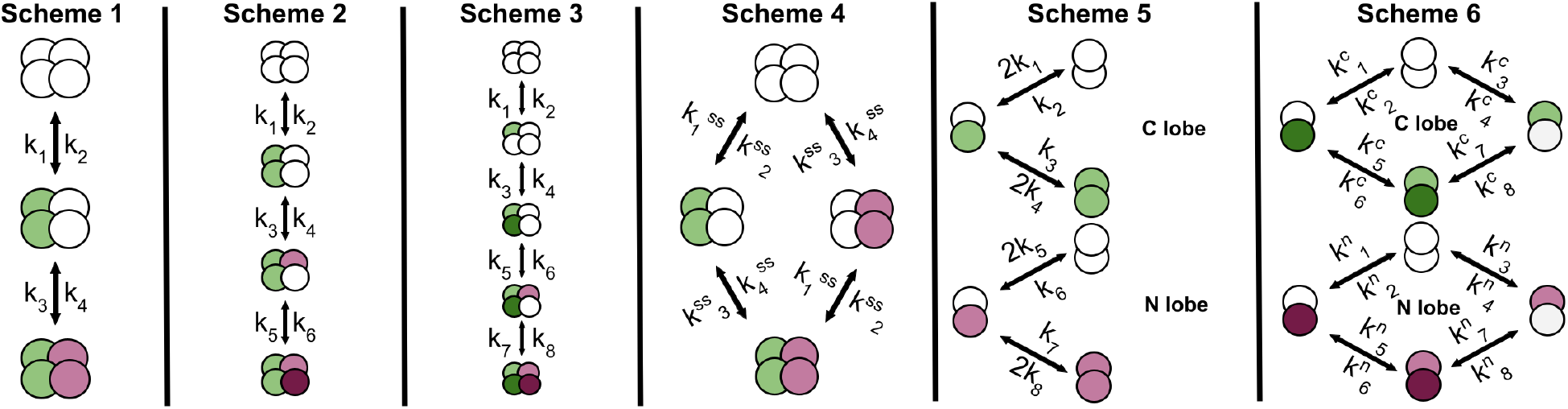
Six calmodulin kinetic schemes to which we fit parameters, and compare to performance with published parameter values. **Scheme 1** – due to strong co-operativity, each calmodulin lobe binds two Ca^2+^ ions at a time, with the lobes modelled sequentially (first C lobe then N lobe). Scheme 1 is parameterised by reaction rate constants 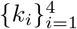. **Scheme 2** – due to co-operativity the first reaction has two Ca^2+^ ions binding as a single event and then the next two Ca^2+^ ions binding sequentially. It is parameterised by reaction rate constants 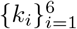 . **Scheme 3** – fully expanded sequential calmodulin scheme where each binding event is represented individually. Depending on the reaction rates, the binding events could be mixed between the lobes, e.g. first binding event could be in the C lobe, the second in the N lobe, or partial combinations of different lobes. The visualised scenario is where the first two events are in the C lobe. This scheme is parameterised by reaction rate constants 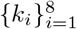. **Scheme 4** – calmodulin binds two Ca^2+^ ions at a time and, contrary to **Schemes 1–3**, the lobes are independent. It is parameterised by eight reaction rate constants 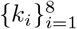 which, along with free Ca^2+^, are used to calculate the effective reaction rates 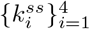 (see **??** for more details). **Scheme 5** – this scheme has independent N and C lobes, with a single Ca^2+^ ion binding at a time. Binding sites within a single lobe are identical. It is parameterised by reaction rate constants 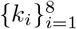 . **Scheme 6** – this scheme has independent N and C lobes, with a single Ca^2+^ ion binding at a time. In contrast to Scheme 5, the binding sites within a single lobe are distinct (indicated by different shades of green/purple). The scheme is parameterised by 16 reaction rate constants 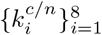 . In all schemes green circles indicate Ca^2+^-bound C lobe sites, purple circles indicate Ca^2+^-bound N lobe sites and arrows indicate bidirectional reactions (Ca^2+^ ions not shown).

The simplest scheme (Scheme 1, Fig 3), from Kim et al. [9], is made up of three calmodulin states – CaM0, CaM2Ca, CaM4Ca, respectively calmodulin bound to no, two and four Ca^2+^ ions. In Scheme 1, Ca^2+^ binding is assumed to be highly co-operative and binding of two Ca^2+^ ions is treated as a single reaction. In principle Scheme 1 does not assume which lobe binds first; the first two Ca^2+^ ions could bind to the C lobe or the N lobe. However, the parameterisation of Scheme 1 by Kim et al. [9] implies that they treat the first Ca^2+^ binding event as being to the C lobe. The published parameter rates in Kim et al. [9] are based on stopped-flow fluorescence measurements [7]. Scheme 1 is parameterised by four reaction rate constants 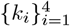 , which can be used to derive two dissociation constants: 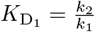 and 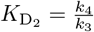.

The next scheme (Scheme 2, Fig 3), from Bhalla and Iyengar [10], is made up of four calmodulin states – CaM0, CaM2Ca, CaM3Ca, CaM4Ca, respectively calmodulin bound to no, two, three and four Ca^2+^ ions. In Scheme 2, binding of the first two Ca^2+^ ions is assumed to be highly co-operative and treated as a single reaction, whereas the next two Ca^2+^ ions bind individually. The parameters in Bhalla and Iyengar [10] (as given in https://doqcs.ncbs.res.in/, also see [28]) do not match neatly to either lobe and the description of how the rates were derived was unavailable at the time of writing. Scheme 2 is parameterised by six reaction rate constants 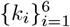 , which can be used to derive three dissociation constants: 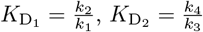 and 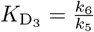.

The final linear scheme that ignores calmodulin lobe-based structure (Scheme 3, Fig 3) is from Shifman et al. [11]. It comprises five calmodulin states – CaM0, CaM1Ca, CaM2Ca, CaM3Ca, CaM4Ca – respectively calmodulin bound to no, one, two, three and four Ca^2+^ ions. The dissociation constants based on experiments in Shifman et al. [11] are 7.9*μ*M, 1.7*μ*M, 35*μ*M, 8.9*μ*M respectively for Ca^2+^ binding events one to four. Reactions in this scheme do not neatly map onto individual Ca^2+^ binding sites within calmodulin lobes; instead they are abstract binding events where, depending on the parameters, they may be probabilistic combinations between different binding sites. Scheme 3 is parameterised by eight reaction rate constants 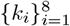 , which can be used to derive four dissociation constants 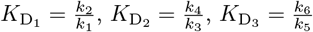 and 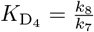. This scheme is the most complex linear CaM scheme possible (without adding conformational calmodulin changes), modelling each Ca^2+^ binding site individually.

Our Scheme 4 (Fig 3) is from Pepke et al. [12] and comprises four states – CaM0, CaM2C, CaM2N, CaM4Ca – respectively calmodulin bound to no Ca^2+^ ions, two at the C lobe, two at the N lobe and four across both lobes. It is the simplest scheme that captures the lobe-based structure of calmodulin. It has eight reaction rate constants 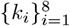 and is based on Scheme 5 (described below), but was simplified used a quasi-steady state approximation for calmodulin species that have a single bound Ca^2+^ ion. This approximation results in elimination of partially bound species from simulations by setting their derivatives to 0 and expressing the partially bound species in terms of the unbound and the fully bound species and permitting the appropriate substitutions in the equations for the unbound and the fully bound species (see **??**).

We draw our Scheme 5 (Fig 3) from model 1 in Pepke et al. [12], which is identical to the scheme used in [5]. It is made up of nine states – CaM0, CaM1C, CaM1N, CaM2C, CaM2N, CaM1C1N, CaM2C1N, CaM1C2N, CaM4 – with the number of Ca^2+^ ions bound to calmodulin indicated by numbers preceding C and N. Even though in total there are nine states, since in this study calmodulin does not bind to any downstream species, we do not need to track individual calmodulin molecules. Therefore, we simulate the lobes as independent species which reduces the number of states from nine to six – CaM0N, CaM1N, CaM2N, CaM0C, CaM1C, CaM2C. Scheme 5 is parameterised by eight reaction rate constants 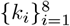, which can be used to derive four dissociation constants 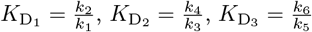 and 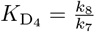 . Pepke et al. [12] used two data sources on calmodulin equilibrium behavior: (1) data from wild-type and tryptic calmodulin fragments (one lobe expressed, other eliminated) [6]; (2) data from competition assays (calmodulin, either wild type or mutants with one active and one inactive lobe, and fluorescent indicator Fluo4FF) [11]. Pepke et al. ( [12], supplemental information) give reaction rate constants as ranges – we take specific numerical values from this model’s entry (model identifier: MODEL1001150000) in the BioModels Database [29, 30]. Faas et al. [5] tuned the model to their own UV-flash photolysis data.

Finally, our Scheme 6 (Fig 3) is from Byrne et al. [13]. It is made up of sixteen states, but similar to Scheme 5, we simulate the lobes as independent species which reduces the number of states to eight – CaM0N, CaMN_1_, CaMN_2_, CaM2N, CaM0C, CaMC_1_, CaMC_2_, CaM2C, where CaM0X denotes an unbound calmodulin lobe, CaMX_1_ denotes Ca^2+^ bound to the first site of a lobe, CaMX_2_ denotes Ca^2+^ bound to the second site of a lobe and CaM2X denotes a fully bound lobe. This scheme is parameterised by 16 reaction rate constants 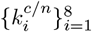 , which can be used to derive eight dissociation constants 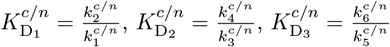 and 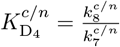 Reaction rates in [13] are based on stopped-flow fluorescence and competitive binding assay data [31].

For each of the six schemes we use the reaction rate constants from the associated publication (we use both Pepke et al. [12] and Faas et al. [5] for Scheme 5). All of the reaction rate constants we used are given in S5 Appendix.

### Ca^2+^ uncaging model

Faas et al. [5] used a linear model for laser induced Ca^2+^ uncaging

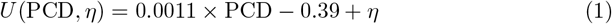

where *U* is the uncaged DMn fraction, PCD is the specified Pockels cell delay, where a larger value results in a higher energy laser pulse and more Ca^2+^ uncaging, and *η* is used to account for the uncertainty of the actual PCD value, i.e. the difference between the specified and physically realised values.

Even though Faas et al. [5] used the linear model successfully in their study, its performance is not quantified and it has some limitations. Most importantly, *U* is not bounded to [0, 1] and can take negative values or values above 1, which would result in physically unrealistic initial conditions. Moreover, it is not clear that a simple linear relationship is optimal to accurately model the relationship between the PCD value and the uncaging fraction. Finally, the variable *η* is additive, and it is not clear that this formulation is optimal – it could be multiplicative or some more complex functional relationship.

Due to lack of the necessary data, we could not develop our own model of how the fraction of calcium uncaged depends on the PCD. Instead, of the linear model (Equation 1) we use an even simpler model that practically performed better than either the linear model from Faas et al. [5] or a neural network. Our uncaging model does not take the theoretical PCD value and contains two learnable parameters, *μ* and Ω for the Gaussian prior distribution of *η* where then *η* is simply passed through a sigmoid function to bound it to [0, 1].

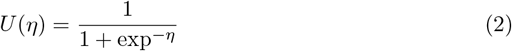

### Fitting our reaction rate constants

We use the `Pumas.jl` [24] Julia package to fit the reaction rate constants, the *η*s which account for the uncertainty of the DMn uncaging fraction (both estimates for individual recordings and variance Ω for their distribution) as well as the observation noise standard deviation *σ*. `Pumas.jl` contains efficient and powerful algorithms for non-linear mixed effects (NLME) modelling, which was essential when fitting the *η*s used to model the uncaging fraction.

Here we adapt the definition provided in [24] to our context, using Scheme 1 as an example. The basis of NLME (shown visually in Fig 4) is the two level hierarchical structure with fixed effects *θ* (reaction rates constants 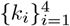 , *μ* and Ω used to sample *η*_*n*_, and observation model noise parameters *σ*) that do not vary between recordings and random effects (*η*_*n*_ used in determining the uncaging fraction) which are individualised for the *n*th recording. Moreover, there is a set of covariates *Z*_*n*_ associated with each recording, i.e. the total concentration of calmodulin, Ca^2+^and OGB5N, which are known. These three sets of values are collated via the structural model *g* into the dynamical parameter vector *p*_*n*_ of the *n*th recording

**Fig 4.**
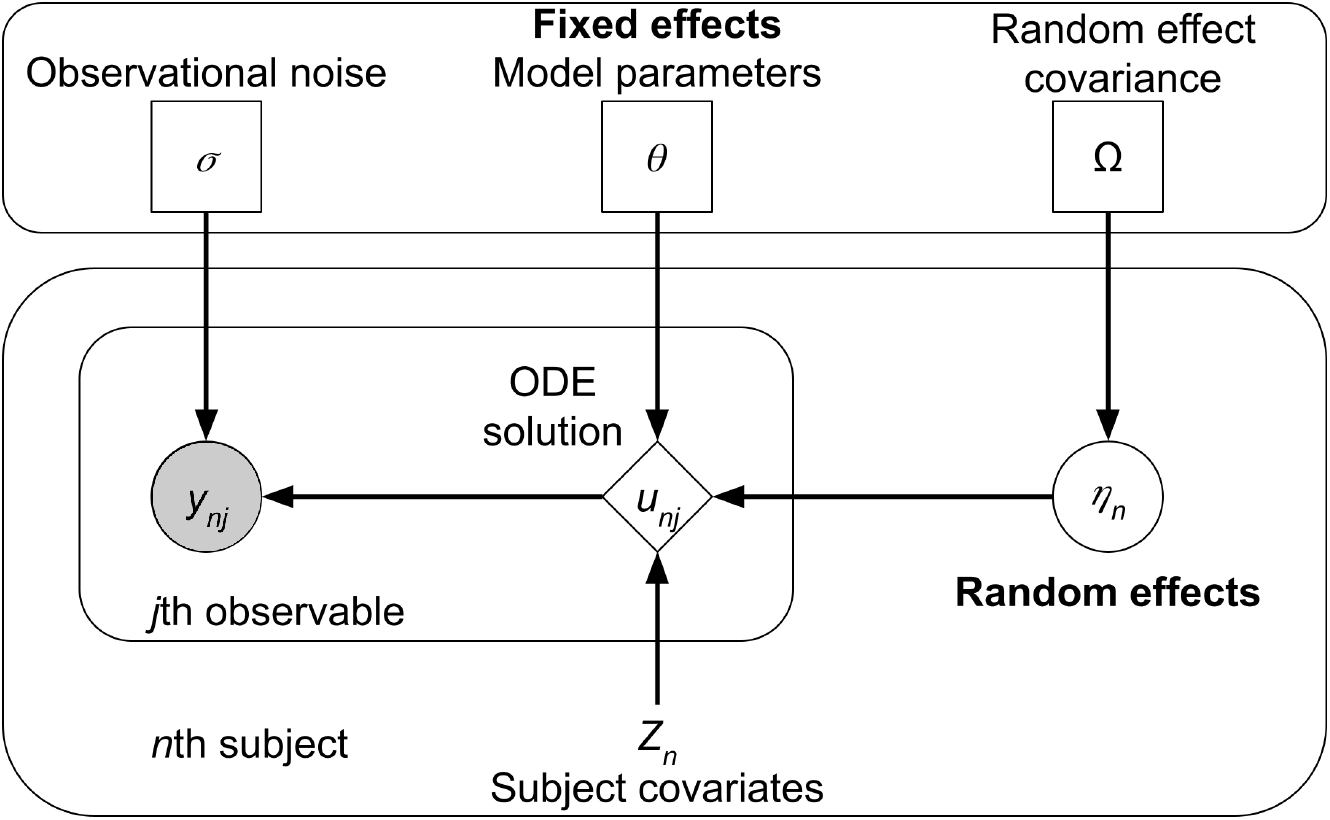
Visual representation of an NLME model, rectangle nodes in the top box denote parameters (fixed effects), circles denote random quantities which are either latent (unfilled) or observed (filled), diamonds are deterministic given the inputs, and nodes without a border are constant.

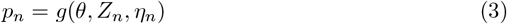

The dynamical parameters *p*_*n*_ are then fed into the model ordinary differential equation (ODE) system

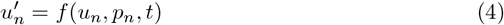

where *u* are the dynamical variables being solved for (DMn, OGB5N, Ca^2+^ and their combinations and the various calmodulin species determined by the scheme being used). For Scheme 1 the system of equations (omitting DMn, OGB5N and Ca^2+^ for brevity) would be

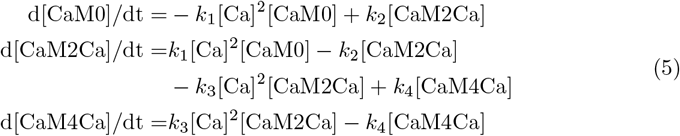

Note that 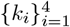 enter into equation above via *p*_*n*_, which can also be used to initialize the ODE system.

The next step is to link the numerical solution of the ODE system to the experimentally observed quantities. The *j*th observable quantity for the *n*th recording at time *t*_*m*_, calculated using *u*_*n*_(*t* = *t*_*m*_) is denoted *y*_*nj*_(*t* = *t*_*m*_) and is obtained through a function *h*_*j*_

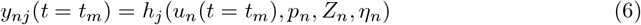

In this study there are two observable quantities: ∆*F/F*_0_ over time being fit to recordings from Faas et al. [5] and Ca^2+^ per calmodulin at equilibrium being fit to Shifman et al. [11]. ∆*F/F*_0_ is derived from OGB5N as follows

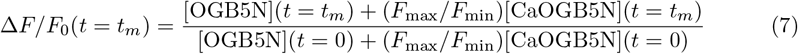

where *F*_max_*/F*_min_ = 39.364 [5]. Ca^2+^ per calmodulin is simply the sum of calmodulin species multiplied by the number of bound Ca^2+^ ions for each species divided by total calmodulin. After obtaining the observable quantities a Gaussian observation model is used to account for observational noise.

There are many ways to fit NLME models, both frequentist and Bayesian [32]. In this study we use the conditional log-likelihood ℒ which for the *j*th observable of the *n*th recording is defined as

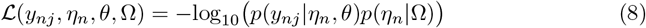

Conditional likelihood is much more numerically efficient compared to other approaches (e.g. marginal likelihood is a two level optimization scheme, Markov Chain Monte Carlo requires many more likelihood evaluations due to sampling). However, conditional likelihood requires appropriate constraints on Ω to avoid overly broad random effect distributions which barely penalize extreme *η*_*n*_ values and effectively result in different individual models due to the learning being offloaded mostly to the random effects.

All fitting was done on the JuliaHub (https://juliahub.com/) cloud computing platform using nodes with 8 vCPUs and 64GB of memory. Individual fits took between one to ten minutes, depending on which scheme was used and whether some of the parameters were fixed. All the code that was used to define the models, run the simulations and perform the analysis is accessible at https://github.com/dom-linkevicius/FaasCalmodulin.jl.git.

### Parameter prior distributions

We incorporated the existing knowledge about calmodulin reaction rates for different kinetic schemes via per-scheme prior distributions that depend on the amount data avaialable for each scheme. All of our priors are in log_10_ space as optimizing rates in log-space was more performant.

For Schemes 1 and 2, since they are simplified and contain fewer reaction rate parameters than an actual calmodulin molecule would, mapping from experimental data to reaction rates is difficult. Therefore, we opted to use wide uniform priors that reflects the small amount of available prior information: 𝒰 (2, 9) for the forward reaction rates (corresponds to 10^2^ M^-2^ms^-1^ to 10^9^ M^-2^ms^-1^) and 𝒰 ( −9, −4) for the dissociation constants (corresponds to a range of 1 nM^2^ to 10 *μ*M^2^).

For Scheme 3, which models Ca^2+^ binding events individually, there is a significant amount of prior information. Specifically, we use set priors on the dissociation constants based on Shifman et al. [11]. We used priors of the form 𝒩 (*r*, 1), where *r* is a dissociation constant from Shifman et al. [11] Table 2 in log_10_. Unfortunately, setting a prior that could similarly constrain the forward reaction rates was not possible, therefore we again opted for wide uniform priors that we used for Schemes 1 and 2: 𝒰 (2, 9).

For Schemes 4 and 5, since they share the same set of reaction rate constants, we used the same set of prior distributions. For each forward reaction rate and dissociation constant we used 𝒩 (*r*, 1), where *r* are rates from Faas et al. [5] in log_10_. We chose Faas et al. [5], rather than Pepke et al. [12], because their rates are based on dynamical data and upon initial simulation runs were performing better. Similarly for Schemes 6, we used the same approach, but centered the Gaussian priors on parameters from Byrne et al. [13].

### Numerical ODE solving

We use the Julia programming language for numerical ODE solving both during and outside of parameter fitting. Specifically, we use the `DifferentialEquations.jl` package [33]. We use the `Rodas5P` numerical solver which can handle significant stiffness in the ODE system and which performed the best of the methods tried. We used it with the default settings, except for reducing the absolute error tolerance to `abs_tol=1e-16` since some simulations that contained low concentrations of species suffered from significant errors in the numerical solution.

### Akaike Information Criterion

There are many ways to compare model performance, but for the purposes of this study we use the Akaike information criterion (AIC) [26]. The AIC for model ℳ with parameters *θ* and given data *d* is defined as

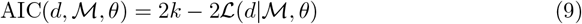

where *k* is the length of the parameter vector *θ*. Therefore, AIC is a measure that evaluates model performance, but also penalizes model complexity via the 2*k* term. There are many other model comparison metrics [34], but the AIC is sufficient for the present study due to the inclusion of predictive model performance and penalizing model complexity along with it being computationally simple to calculate.

## Results

### General model fitting results

We fit each of the six kinetic schemes shown in Fig 3 to the fluorescence traces from Faas et al. [5] and the steady state calcium-calmodulin binding data from Shifman et al. [11]. We used the root mean square error (RMSE) to evaluate the goodness of fit between the models and the data. The fitting procedure was repeated 20 times with different random seeds, which set the random sampling of the training, validation and test data (see S1 Appendix) along with the initial parameters for optimization. Therefore, due to variability in training data and initial parameters, RMSE values (especially for our parameter fits) for each seed can be significantly different.

We now compare the performance of each kinetic scheme with the reaction rate constants we fit and the published ones. Table 1 shows the training (split between dynamical data from Faas et al. [5] and equilibrium data from Shifman et al. [11]), validation and test data set performance summary statistics for the six investigated kinetic schemes, reporting the average RMSE values for the 20 different seeds used. The distribution of the test RMSE values is shown in Fig 5. On average Schemes 5–6 performed the best, whereas other schemes were not able to capture either the equilibrium data (Schemes 3–4) or both the dynamical and the equilibrium data (Schemes 1–2).

**Table 1.** Summary of training, validation and test performance (RMSE *±* SD) for different kinetic schemes with either parameters fit from scratch, fixed to values from publications or our modifications.

| Model | Training<br>(Dynamical<br>data) | Training<br>(Equilibrium<br>data) | Validation<br>(Dynamical<br>data) | Testing<br>(Dynamical<br>data) | Cohen’s d<sup>‡</sup> |
| --- | --- | --- | --- | --- | --- |
| **Scheme 1 + our rates** | **0.74 ± 0.12** | **1.47 ± 0.17** | **0.80 ± 0.16** | **0.77 ± 0.17** | - |
| Scheme 1 + Kim et al. rates | 1.51 ± 0.20 | 2.31 | 1.16 ± 0.06 | 1.11 ± 0.08 * | 2.07 |
| **Scheme 2 + our rates** | **0.72 ± 0.09** | **2.00 ± 0.28** | **0.85 ± 0.15** | **0.81 ± 0.17** | - |
| Scheme 2 + Bhalla and Iyengar rates | 1.03 ± 0.04 | 2.23 | 1.16 ± 0.06 | 1.11 ± 0.08 * | 1.77 |
| **Scheme 3 + our rates** | **0.43 ± 0.03** | **0.83 ± 0.46** | **0.48 ± 0.05** | **0.46 ± 0.05** | - |
| Scheme 3 + Shifman et al. rates<sup>†</sup> | 0.61 ± 0.05 | 0.46 | 0.70 ± 0.11 | 0.66 ± 0.10 * | 3.76 |
| **Scheme 4 + our rates** | **0.42 ± 0.06** | **0.88 ± 0.10** | **0.46 ± 0.12** | **0.44 ± 0.11** | - |
| Scheme 4 + Pepke et al. rates | 0.82 ± 0.05 | 0.78 | 0.84 ± 0.06 | 0.82 ± 0.09 ** | 3.47 |
| **Scheme 5 + our rates** | **0.37 ± 0.02** | **0.44 ± 0.18** | **0.40 ± 0.03** | **0.38 ± 0.04** | - |
| Scheme 5 + Faas et al. rates | 0.45 ± 0.02 | 0.75 | 0.56 ± 0.03 | 0.53 ± 0.03 ** | 4.17 |
| Scheme 5 + Pepke et al. rates | 0.86 ± 0.05 | 0.75 | 0.88 ± 0.07 | 0.86 ± 0.09 ** | 13.5 |
| **Scheme 6 + our rates** | **0.35 ± 0.02** | **0.35 ± 0.03** | **0.38 ± 0.03** | **0.36 ± 0.03** | - |
| Scheme 6 + Byrne et al. rates | 0.47 ± 0.01 | 0.83 | 0.55 ± 0.03 | 0.53 ± 0.02 ** | 5.83 |

Highlighted rows were the best performing for that scheme. Note that for models with fixed reaction rates trained on Shifman et al. [11] data, no SD is given since all of the data is used for training and there is no variance in this metric. T-tests are done with reference to our reaction rate constants.
<sup>†</sup> Shifman et al. [11] only contained dissociation constant values, therefore we had to fit on-rates while fixing to their $K_D$ values.
<sup>‡</sup> Effect sizes evaluated via Cohen’s d over 1.2 are considered very large and over 2.0 are considered huge [35].
\* $p < 10^{-6}$
\*\* $p < 10^{-10}$

**Table 2.** Median AIC values for all combinations of kinetic schemes and reaction rates (our fits or published).

| | Median AIC values ($\times 10^3$) | | | | | |
| --- | --- | --- | --- | --- | --- | --- |
| Parameter source | Scheme 1 | Scheme 2 | Scheme 3 | Scheme 4 | Scheme 5 | Scheme 6 |
| Ours | 12.9 | 14.9 | 8.32 | 6.37 | 6.45 | 5.97 |
| Published | 17.9 | 18.4 | 11.5 | 14.2 | 10.2<sup>†</sup> / 14.8<sup>‡</sup> | 9.21 |

<sup>†</sup> when rates from Faas et al. [5] are used.
<sup>‡</sup> when rates from Pepke et al. [12] are used.

**Fig 5.**
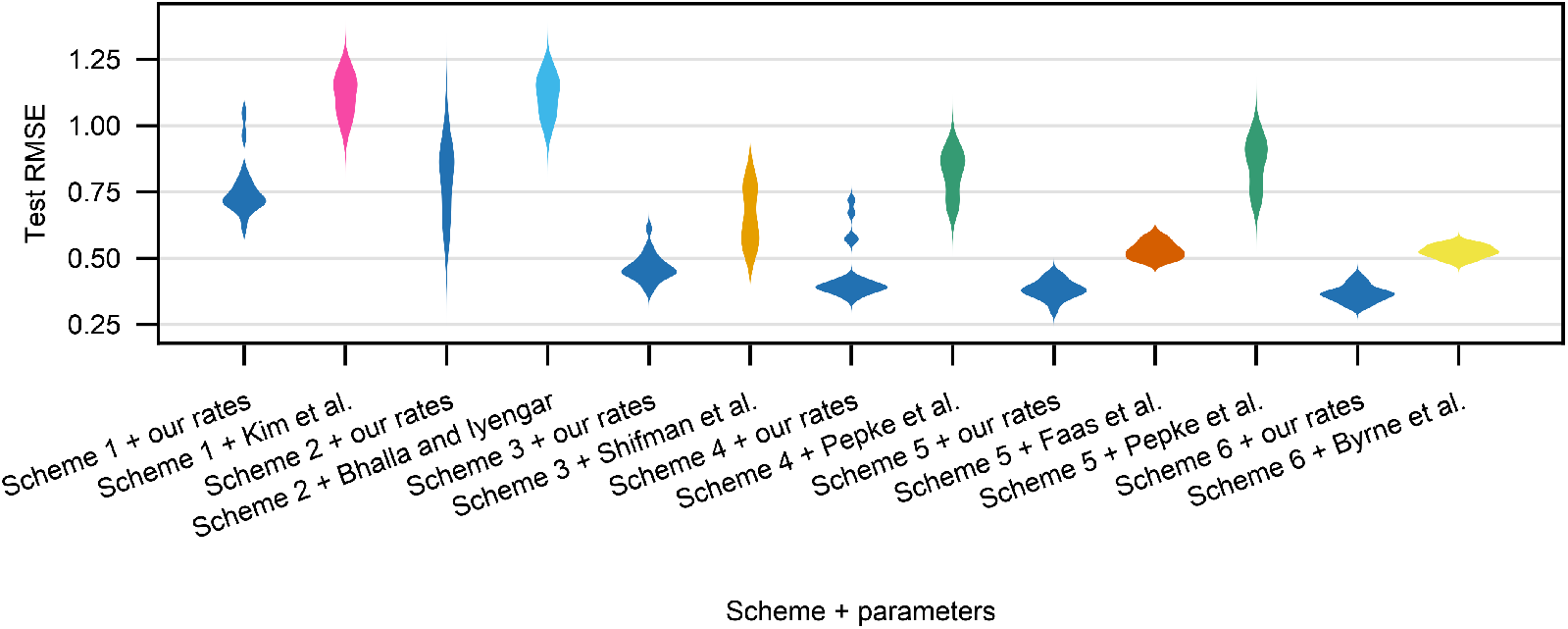
Violin plots of RMSE values for the test data set for each seed for all schemes for our own and the published parameter sets.

#### Dynamical behaviour

To illustrate the differences between model fits, Fig 6 shows the measured fluorescence traces that were used as the validation data set for seed 1 and corresponding model predictions for each kinetic scheme with our fitted rates and the published rates. Each row shows the same 24 experimental traces (black), split between the seven groups of solutions which were used experimentally in Faas et al. [5]. Within each group, each trace corresponds to a different laser uncaging strength used – the stronger the laser, the more calcium gets released, the larger the ∆*F/F*_0_ values that are measured.

**Fig 6.**
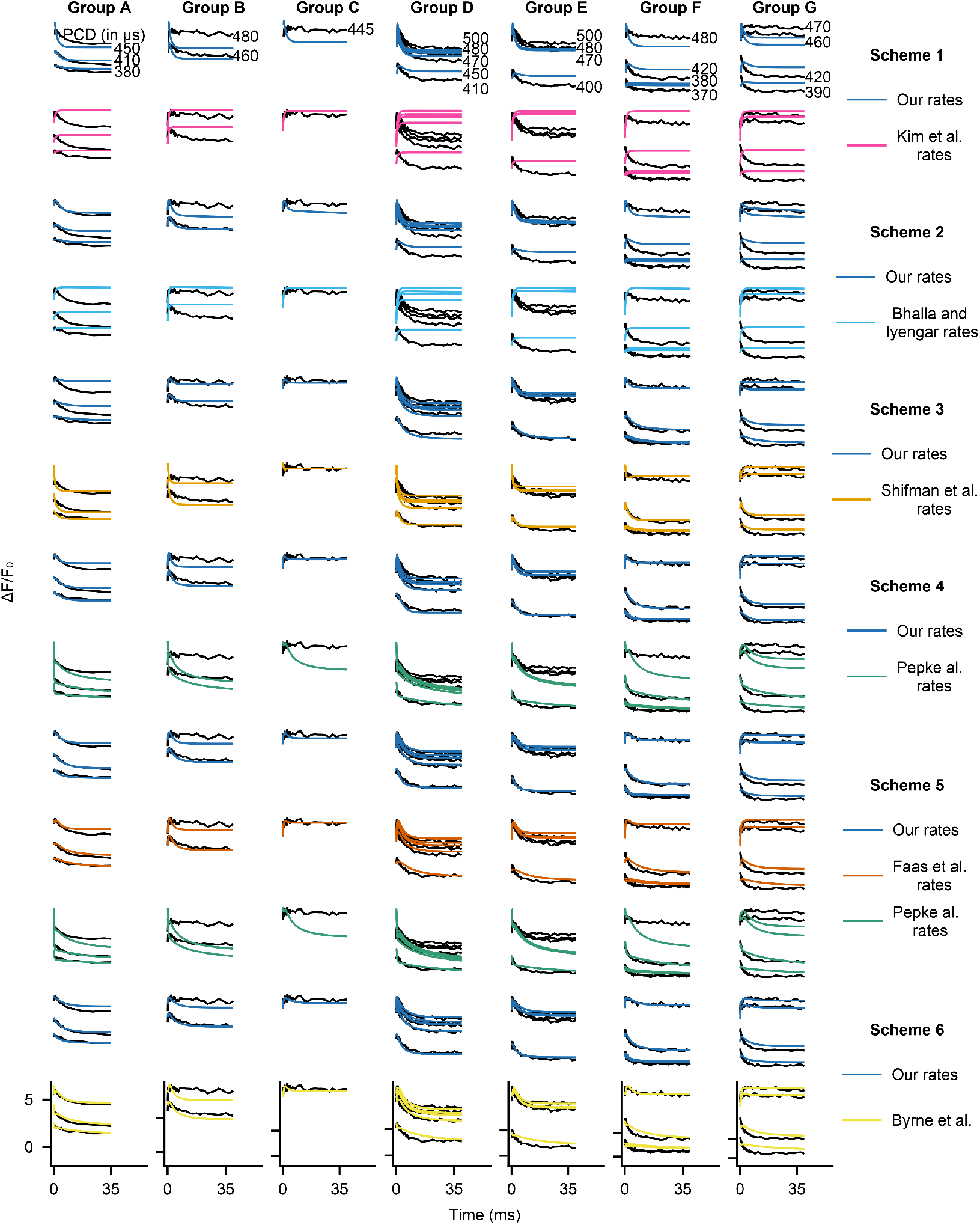
Sample dynamics for validation data for seed 1 for all trained models. Each column is a single data group A–G from Faas et al. [5], whereas each row is a different combination of scheme + reaction rate constants (specific combination given in the rightmost column). In all subplots y axis is ∆*F/F*_0_ and x axis is time. Black lines are empirical data and red lines are model outputs. Scale bars are given at the bottom row, each tick on the y-axis corresponds to the same value in each column.

The biggest differences in measured and predicted dynamics are for Scheme 1, for which the RMSE differences are also the largest. In Fig 6 the main difference between our parameters and rates from Kim et al. [9] is that our rates give rise to traces that follow the experimental dynamics to some extent, whereas the published rates simply equilibrate to a value and barely display any dynamics (e.g. groups D–G). However, even though our rates result in dynamical behaviour, they do not show good equilibrium performance and in fact do not reach equilibrium when the data has long reached it.

The comparison of dynamics for Scheme 2 is similar to that of Scheme 1. Comparing our reaction rates with those in Bhalla and Iyengar [10], we see that the published reaction rates make the system equilibrate and not follow the data closely, whereas our reaction rates achieve a more accurate fit. However, with either our rates or the published rates, the traces and the average RMSE values indicate that Scheme 2 fits the data quite poorly.

Models with the level of complexity of Scheme 3 and higher are able to capture the dynamical data much better than the simpler Schemes 1 and 2. As shown in Fig 6, both Scheme 3 models perform adequately. However, our reaction rates trained from scratch still perform better, especially in capturing the initial rise and fall in ∆*F/F*_0_ (for example see columns D–F). Note that for this scheme the comparison is not entirely equivalent to other cases as we had to fit the forward reaction rates while we kept the dissociation rates fixed to those in Shifman et al. [11].

For Scheme 4, our reaction rate constants significantly outperform those of Pepke et al. [12] as shown in Fig 6. This is reflected in a smaller mean RMSE value of our rates and is evident in most experimental groups, where our rates result in reasonably accurate predictions whereas the Pepke et al. rates significantly under-predict ∆*F/F*_0_.

Looking at the dynamics for Scheme 5, based on the RMSE values in Table 1, our rates perform significantly better than the rates from either Faas et al. [5] or Pepke et al. [12], but the gap is much smaller for the former than the latter. The differences in dynamics between our fits and rates in Faas et al. are subtle, but generally our rates perform better for small amounts of uncaged Ca^2+^. In contrast, comparing dynamics with our rates to dynamics with rates in Pepke et al. [12], their reaction rates result in significant mismatches to the data, to the point that the optimization procedure has to inject amounts of Ca^2+^ that lead to incorrect equilibrium levels (see Fig 7, which shows that their dissociation constants can fit equilibrium data well).

**Fig 7.**
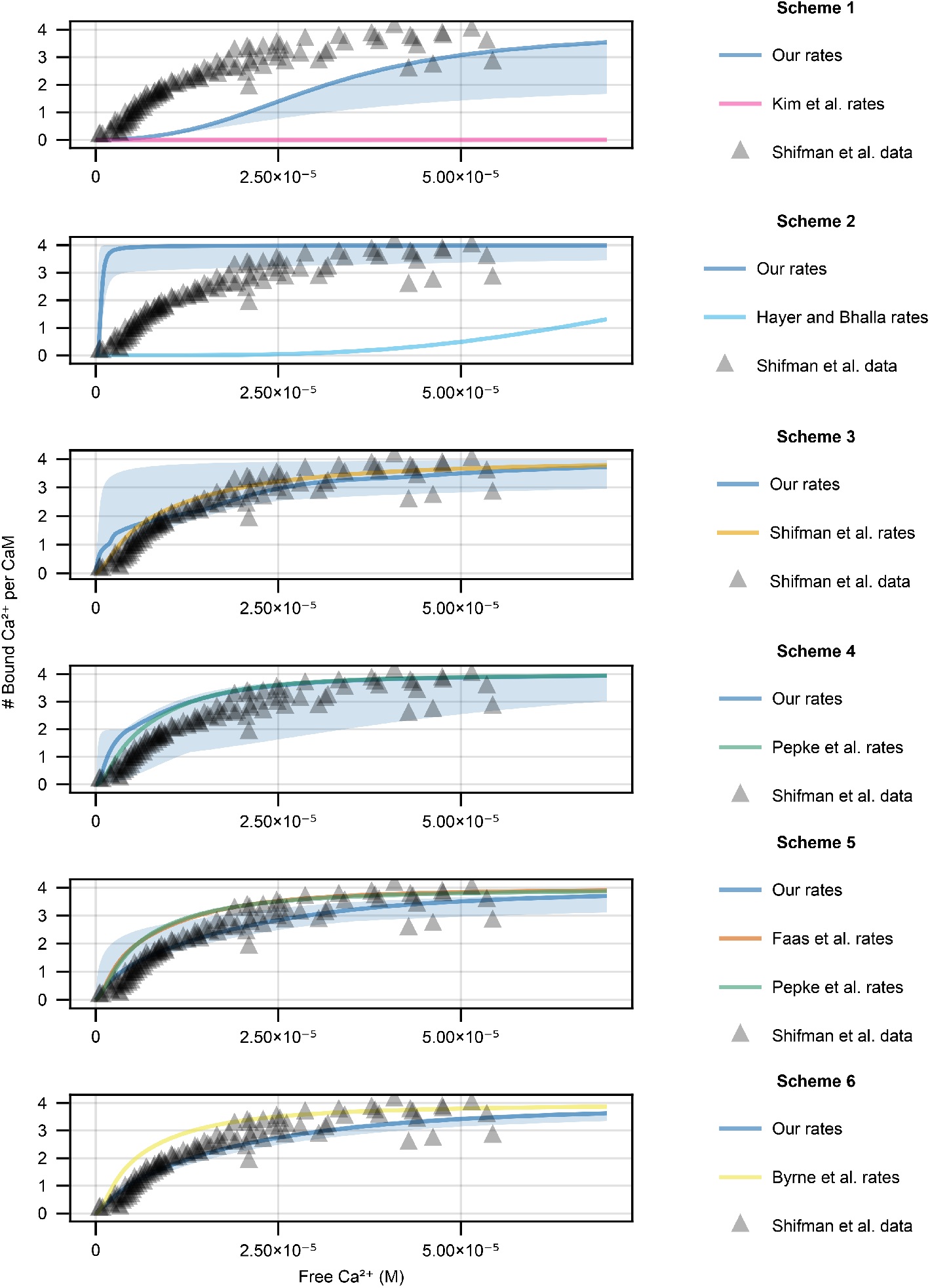
Calmodulin equilibrium behaviour when the free amount of Ca^2+^ is varied for all kinetic schemes for the 20 seeds that we tested (solid line is the median and shaded area is the 95% confidence interval). Model behaviour with our reaction rate constants are plotted in blue, whereas published ones are in the colors indicated. Note that at times the published reaction rate constants (Scheme 1 and Scheme 2) result in behaviours that are much more right-shifted, therefore show up as zero in the relevant range. We also include data points for wild type calmodulin for an equivalent experimental setup in Shifman et al. [11].

Finally, for Scheme 6, the main differences between the dynamics resulting from our reaction rates and those in Byrne et al. [13] are generally seen for small amounts of uncaged calcium (columns D–G bottom traces). Our reaction rates (for this seed) managed to capture calmodulin behaviour with low amounts of Ca^2+^ more accurately. Even though for some traces the published rates can outperform ours (e.g. top traces in either groups A or C), our reaction rates on average show a smaller RMSE value.

#### Equilibrium behaviour

Fig 7 shows the equilibrium behaviours of our reaction rate constants (for all 20 training seeds) and published ones. When using Scheme 1 (Fig 7, top row), the fits to data from Shifman et al. [11] are generally poor. A single run was able to fit the equilibrium data well, but in most fits, calmodulin was less sensitive to Ca^2+^ than indicated by the data. Some of this behaviour can be attributed to the prior because most of the runs hit the uniform distribution limits (especially the dissociation constants). When the limits were wider, there was a significant amount of training failures (up to 30%) and trained models showed step-like equilibrium behaviour that was overly sensitive to Ca^2+^.

Therefore, we opted to keep the narrower limits. Scheme 1 with our parameter sets is generally much more sensitive to Ca^2+^than it is with the rates from Kim et al. [9], which result in calmodulin behaviour that does not show appreciable Ca^2+^ binding in the relevant Ca^2+^ range and is significantly right-shifted (Fig 7, top row pink line).

For Scheme 2 (Fig 7 second row from the top), our fits result in behaviour that is significantly more sensitive to Ca^2+^ than the experimental data. Calmodulin would be close to fully bound under resting neuronal Ca^2+^ levels, in contrast to the reaction rates from Bhalla and Iyengar [10], which are significantly less sensitive to Ca^2+^ than the data indicate, not reaching full calmodulin saturation in the experimental data range. The failure to fit the equilibrium data is likely due to the inclusion of the dynamical data – reaction rate sets that would allow this scheme to fit equilibrium data do not fit the dynamical data well. The failure of Scheme 2 fitting both dynamical and equilibrium data is likely due to the first Ca^2+^ binding event including two Ca^2+^ ions and needing to be relatively fast to fit the dynamical data.

For Scheme 3 (Fig 7 third row from the top), the parameters from Shifman et al. [11] perform very well because they were explicitly tuned to only this data set. However, when the dynamical data from Faas et al. [5] is included in the fitting procedure, the resulting equilibrium behaviour varies between runs (blue shaded area). Similarly to Scheme 2, our fits result in behaviour that is much more sensitive to Ca^2+^ than the data indicates. However, contrary to Scheme 2, the range of behaviours is much more varied and a significant portion of fits match data from Shifman et al. [11] reasonably well.

For Scheme 4 (Fig 7, fourth row from the top), even though in general the fits are much better, there are still a few runs that do not perform as well. Moreover, our mean RMSE value is slightly worse than that parameters from Pepke et al. [12] for the equilibrium data in Shifman et al. [11]. Curiously the mean RMSE for the dynamics predicted via Scheme 4 with our rates is much smaller compared to the rates from

Pepke et al. [12]. A possible explanation for this is that rates from Pepke et al. were first derived using Scheme 5 and then reduced to Scheme 4. Scheme 5 is a more powerful model due to having more state variables which could make optimization easier than simply using Scheme 4 (see results below).

Our fits to the Shifman et al. [11] equilibrium data follow a similar pattern for Schemes 5 and 6. For both Schemes the noisiness in model behaviour that was present for Schemes 1–4 is either gone or significantly smaller, and most fits match the data from Shifman et al. [11] reasonably well. In both cases, the average RMSE value using our rates is significantly smaller compared to the published reaction rates.

### Model comparison via AIC

We now compare both the published reaction rates to our reaction rates and between the kinetic schemes via AIC evaluated on the test set of a random seed. AIC is a useful model comparison tool because it takes into account both model predictive performance as well as model complexity (number of parameters). Fig 8 shows the box plots for the AIC values for all the combinations of kinetic scheme and parameter set for all 20 seeds. As shown in Table 2, our reaction rate constants have lower median AIC values (lose less information) compared to the published ones. Moreover, the more complex the scheme, the lower the AIC value, with Scheme 6 performing the best. The median AIC values seem to asymptote and reach a lower value by Scheme 6 (the big change is after Schemes 2–3), so only qualitatively different model improvements are likely to decrease the AIC value more.

**Fig 8.**
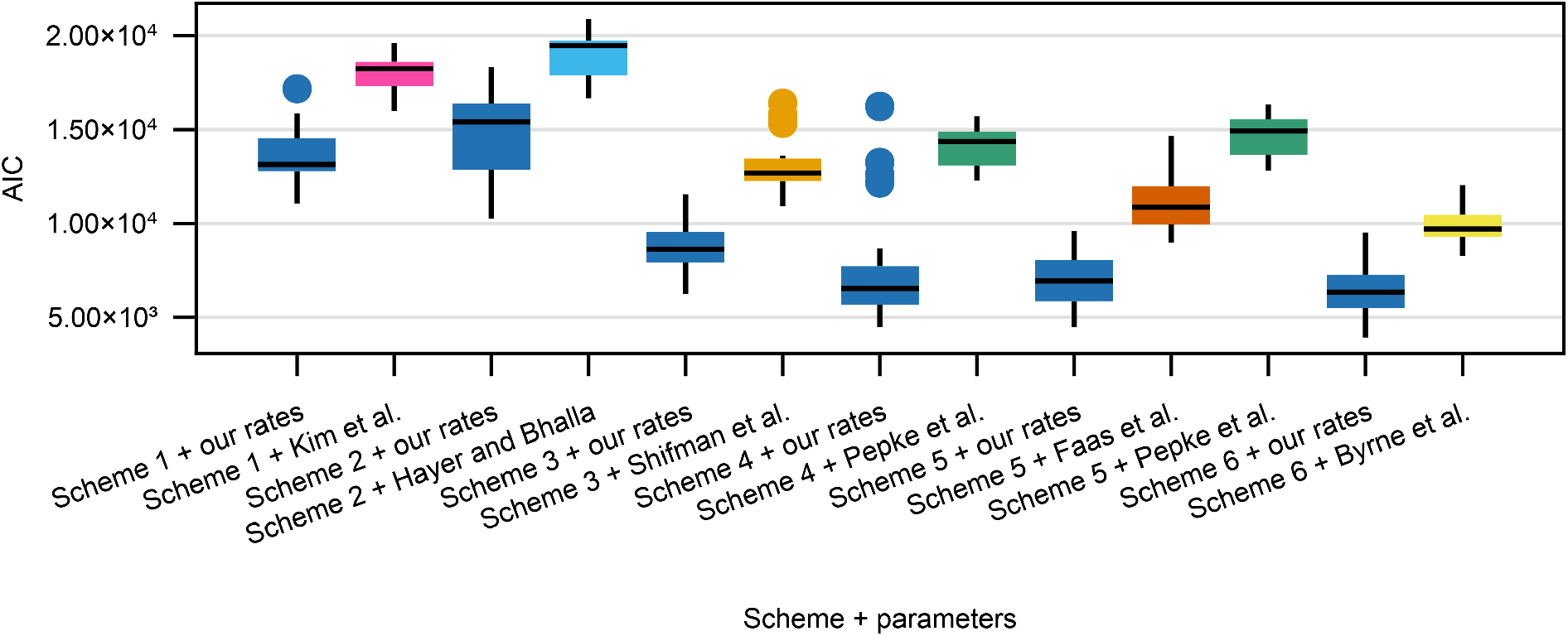
AIC box plots for all the different combinations of kinetic schemes and either previously published or our own reaction rate constants. Each box plot is based on training the model on 20 different random seeds, the AIC value is calculated on the test data set of a given seed.

Calculating the relative likelihoods from median AIC values, where our reaction rate constants are the reference, all the published parameter sets have negligibly low relative likelihoods (largest being on the order of *e*^−100^). Therefore, our reaction rate constants are significantly more likely compared to the published ones.

Since our reaction rate constants have lower AIC values (lose less information) compared to the published ones, we use our reaction rate constants to compare between different kinetic schemes. Given the results in Fig 8 and Table 2 and using the median AIC for Scheme 6 with our rates as reference, the other schemes with our parameter sets have a negligibly small relative likelihoods (again on the order of *e*^−100^). Therefore, of the combinations of schemes + reaction rates that we found, Scheme 6 with our reaction rates is relatively the most likely.

### Calmodulin Ca^2+^ integration properties

Having established that our reaction rate constants are significantly more likely than the published ones, we now ask whether this difference is meaningful practically. To answer this question we probe the Ca^2+^ signal integration properties of calmodulin. CA1 pyramidal cell Schaffer collateral synapses undergo long-term potentiation dependent on CaMKII (and therefore on calmodulin) in response to three 1s trains of 50Hz stimulation [36]. Given these results, it is likely that calmodulin integrates the Ca^2+^ signal within a single train. Therefore, we set up a series of simulations where a model was stimulated by a 1s train of Ca^2+^ injections but the frequency was varied from 2Hz to 100Hz. Based on the results in Sabatini et al. [37], Ca^2+^ influx due to single synaptic stimulation event for a neuron at resting voltage is around 0.7*μ*M (this mimics the experimental setup in Bayazitov et al. [36] best). To mimic the competition between calmodulin and other buffers and pumps we implemented a minimal Ca^2+^ extrusion model using values in Sabatini et al. [37] Table 1 for a CA1 pyramidal cell spine – Ca^2+^ decaying to a baseline of 100nM with a time constant *τ* = 12ms. Finally, we use a biologically realistic calmodulin concentration of 20*μ*M [38]. After the simulation, we evaluate the calmodulin signal integration by calculating the area under the curve of both partially bound calmodulin and fully bound calmodulin, where bigger values indicate a larger level of Ca^2+^ signal integration.

As shown in Fig 9 columns one and two, Schemes 1 and 2 are not capable of integrating Ca^2+^ signals in the tested frequency range (except for a few outlier runs with Scheme 2). For Scheme 1 it is likely due to the fact that the models are not sensitive enough to Ca^2+^ (see Fig 7 top row). The same interpretation, however, does not hold for Scheme 2, whose equilibrium behaviour with our parameter fits was usually too sensitive to Ca^2+^ compared to experimental data. This behaviour for Scheme 2 may be explained by slow reaction rate constants compared to Ca^2+^ decay.

**Fig 9.**
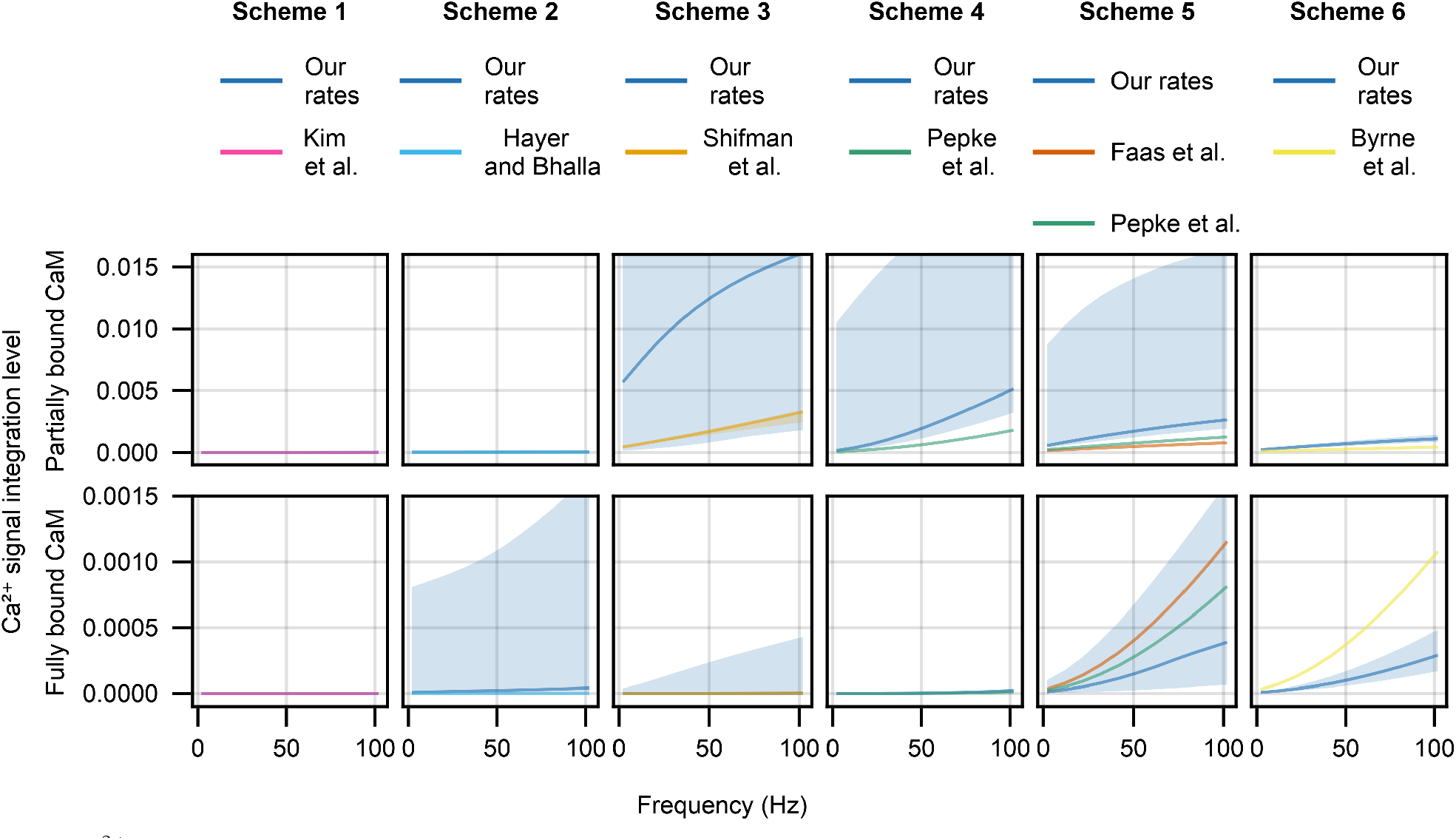
Ca^2+^ signal integration properties (measured as area under the curve) of partially (first row) and fully bound (second row) calmodulin species in response to a 1sec train of 2–100Hz stimulation that delivers 0.7*μ*M Ca^2+^ per spike (same general patterns hold if 12*μ*M Ca^2+^ per spike is delivered, results not shown). A different calmodulin scheme is used in each column and shows our parameter fits (deep blue lines) and the published parameter values (all other colours). The solid lines are median model behaviour and shaded areas are the 95% confidence intervals.

The results are somewhat different for Schemes 3 and 4 (Fig 9 columns three and four), where both the published reaction rates and our own fits show significant Ca^2+^ integration in the partially bound calmodulin species, but barely any in the fully bound calmodulin. In both cases our reaction rates result in significantly higher Ca^2+^ signal integration. However, results for Scheme 3 with our reaction rates which show significant Ca^2+^ integration at 2Hz should be taken with caution due to the same fits being overly sensitive to Ca^2+^ at equilibrium (Fig 7 third row from the top).

Finally, for Schemes 5 and 6 we see integration of Ca^2+^ signals that results in both fully and partially bound calmodulin species (Fig 9 columns five and six). For both schemes our reaction rate constants predict that Ca^2+^ signal integration would result in more partially bound calmodulin compared to predcitions from the published rates. As for fully bound calmodulin, our reaction rates predict a lower level of fully saturated calmodulin than the predcitions from the published rates.

The difference between the partially and fully bound calmodulin signals is more pronounced with our reaction rates than with the published ones. For Scheme 5 the difference is around an order of magnitude for our reaction rates and under 2-fold for the published reaction rates, whereas for Scheme 6 the difference is around 4-fold for our reaction rates and around 2-fold for the published reaction rates. Given that our models reaction rates perform better, we predict that partially bound calmodulin species play a more significant part in Ca^2+^ signalling integration and propagation than predicted by previously published models.

### Parameter Correlations

We next examine the relationships between our parameter fits within a given scheme. Analysing relationships between parameters may point future experimental research questions. For example, if some reaction rate constants are correlated, they may be under-determined. Therefore, future model development would benefit from additional, more directed data to better constrain the correlated parameters. We use partial correlation as a measure of relationship between parameters [19]. Briefly, partial correlation quantifies the degree of association between two variables when the variance from a set of controlling variables is taken into account. For example, in Scheme 1 the partial correlation between *k*_1_ and *k*_3_ would indicate the relationship between these two rates when 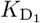 and 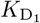 is accounted for.

Since structurally there is nothing to distinguish between the C and the N lobes for Schemes 4–6, we calculate the dissociation constant for the two binding reactions in Scheme 4 and for the first binding reaction for both lobes in Scheme 5 and Scheme 6 and compare their values – the one that has a lower *K*_D_ value we call the C lobe and the one that has a higher value we call the N lobe. This is to avoid what we call the C lobe being functionally the N lobe and vice versa, which would result in artificially higher parameter spread or obscure parameter correlations. We show the parameter pair plots for all six schemes in the S6 Appendix, along with tables of individual reaction rate fits. We show partial correlation coefficients for all schemes in Fig 10.

**Fig 10.**
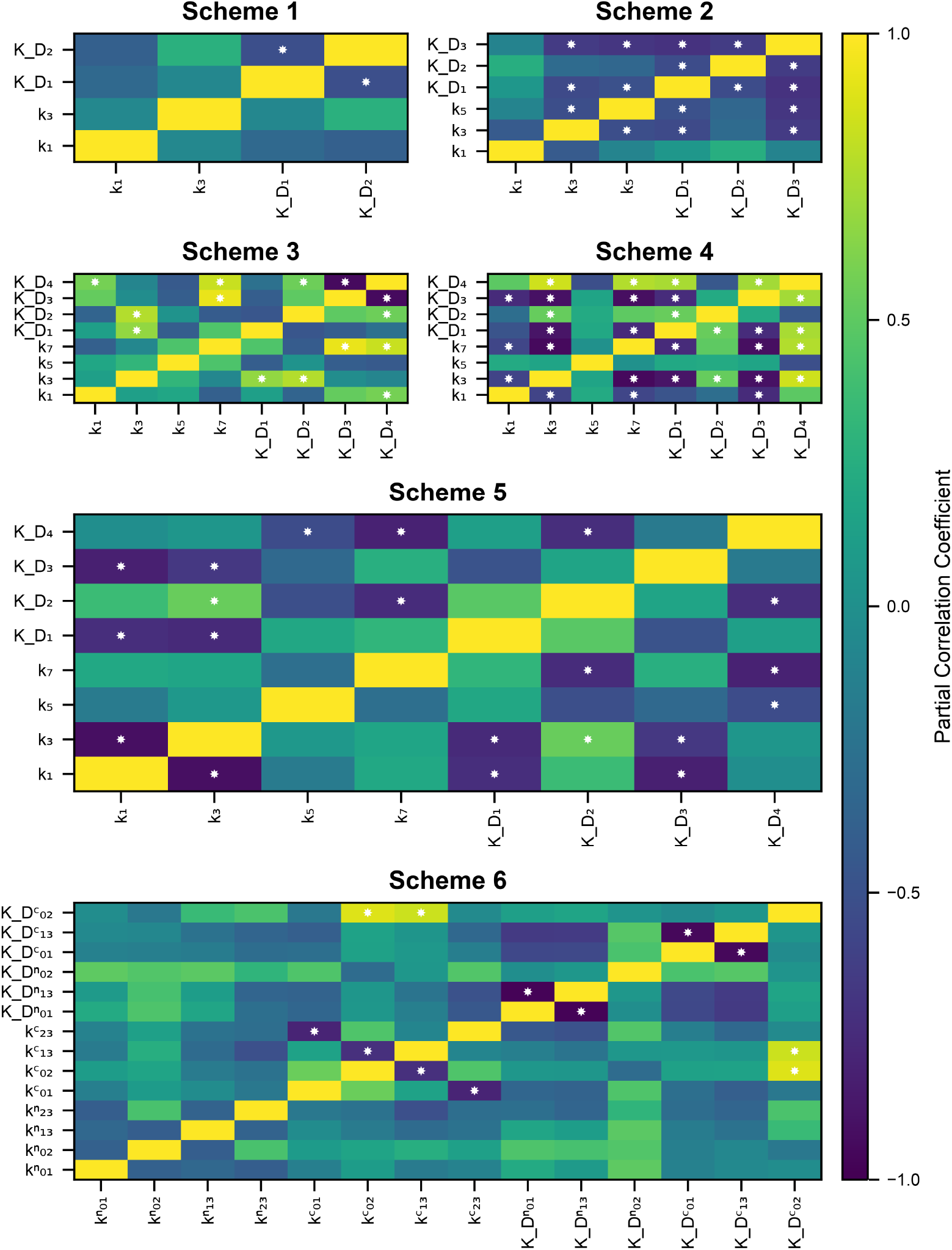
Partial correlation coefficients for our reaction rate constant fits for all schemes. White stars in the rectangle indicate *p <* 0.05 with null hypothesis that the partial correlation is zero.

First of all, even though the data set we use is richer, as it includes both the dynamical and the equilibrium data, there is still significant variance in our model parameter fits. For some parameters the pairs can span 5 to 10 orders of magnitude (see S6 Appendix). Moreover, there are significant correlations between multiple parameters in most schemes.

There is only one significant negative correlation for Scheme 1, between 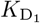 and 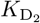 . This correlation is most likely due to a limited number of degrees of freedom offered by this scheme. Assuming that a calmodulin molecule has an overall dissociation constant that is a function of the dissociation constants of individual reactions, the dissociation constants of individual reactions have to co-vary in order to maintain the same overall behaviour.

Reaction rate constants for Schemes 2 have 8/15 significant correlations which indicates a high level of sloppiness in the system. Similarly to Scheme 1, given the high level of simplification used in this scheme, in order to maintain the same overall model behaviour, parameters have to co-vary. This is especially true for the dissociation constants, which are all are negatively correlated. We also see that the on-rate for the first reaction is not correlated to any other parameter and is therefore fairly well-determined.

Scheme 3 has proportionally fewer correlations than Scheme 2 (7 out of 28 parameters correlated). Most correlations are with the dissociation constant for the final Ca^2+^ binding event. This can be explained by thinking about a general dissociation constant for calmodulin, which would be a function of 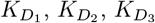 and 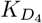 since 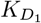is likely better constrained due to the first binding event needing to be of particular speed to fit the rising/falling phase of the individual time series, the other dissociation constants may co-vary more freely and balance each other out. The on-rate for the first binding event is only correlated to one variable, which indicates that it is quite well determined.

Scheme 4 shows the largest fraction of correlations of all the schemes (15/28). This is most likely due to the quasi-steady state approximation which results in steady state reaction rates 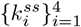 that provide a lot of room for sloppiness via products and quotients of the full set of reaction rates 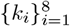 .

The point that model structure is of utmost importance in determining the levels of sloppiness in the system is further reinforced by Scheme 5, where 10 out of 28 reaction rates were correlated. More importantly, a significant number of correlations are within-lobe, for example *k*_1_ and *k*_3_ – the first and second on-rates. There are also some cross-lobe correlations, for example 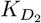 and 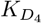 which are the second Ca^2+^ binding events for C and N lobes respectively.

Curiously, even though Scheme 6 is the most complex in terms of number of parameters and number of states, it shows only six significant correlations between reaction rate constants. Moreover, all correlations are within a lobe, rather than between lobes. More specifically, most of them are for parameters in the C lobe, rather than the N lobe.

### Necessary structural components of a calmodulin model

As shown in Table 1 and in Fig 6, there is a large gap in training performance between Schemes 1–4 and Schemes 5–6. Even though training RMSE in both dynamical and equilibrium data significantly decreases going from Scheme 2 to Scheme 3, only from Scheme 5 onwards can both dynamics and equilibrium behaviour be captured well.

There are two main differences between Schemes 1,2,4 and and 5–6: independence of lobes and structural assumption of co-operativity. Both Scheme 3 and Schemes 5–6 allow co-operativity (via reaction rates) but do not assume it structurally. Schemes 3 does not allow for independence of lobes, while Schemes 5–6 assume it structurally. In this section we provide an empirical argument that links model features to gaps in performance, focusing on event-based (as opposed to binding site-based) and structurally co-operative (especially for the C lobe) schemes to model calmodulin.

Assuming that the real calmodulin dynamics operate in a *k*-dimensional space, any model capable of modeling the dynamics would have to have at least that many dimensions (along with an appropriate structure). Calmodulin models framed in terms of events (fully abstracted from binding sites) can operate at most in a four dimensional linear subspace (since rank of such a network is four, see page 30 in [39]) of the five dimensional state space (see Scheme 3 in Fig 3). Therefore, an immediate conclusion of this may be that *k >* 4, real calmodulin dynamics operate in a higher dimensional space than an event-based model allows for. However, Scheme 5, which is able to model both calmodulin dynamics and equilibrium behaviour (see Table 1), has rank 4 as well. The main difference between Schemes 3 and 5 are the independence of the lobes: Scheme 5 contains two independent subnetworks (each of which is rank 2). Therefore, based on our results, in order to accurately model both calmodulin dynamics and equilibrium behaviour, two independent subnetworks (independence of lobes) is a necessary model feature.

We next analyze whether a structural assumption of co-operativity, modelling the binding of two Ca^2+^ ions as a single event, within calmodulin lobes is reasonable. This is not the only way of modelling co-operativity, but it results in models with a smaller state space vector and therefore can be preferable computationally. Fractional calmodulin occupancy of the N and the C lobes using a well performing model (Scheme 6 with parameters from Byrne et al. [13]) is shown in Fig 11 columns one and two. Starting with the dynamics of the partially occupied N lobe, the model predicts around 20% of calmodulin molecules would have the first site occupied, with a negligible fraction having the second site occupied. Moreover, the dynamics of partially occupied sites in the N lobe do not show fast changes over the simulated time period, so the quasi-steady state approximation would hold reasonably well. The dynamics of the C lobe paint an opposite picture. It is immediately obvious that, due to its slower speed, the quasi-steady state approximation (d[CaMC_1_]*/*d*t* = 0) does not hold for the C lobe as there are calmodulin dynamics occurring over the whole simulated time of 35ms.

**Fig 11.**
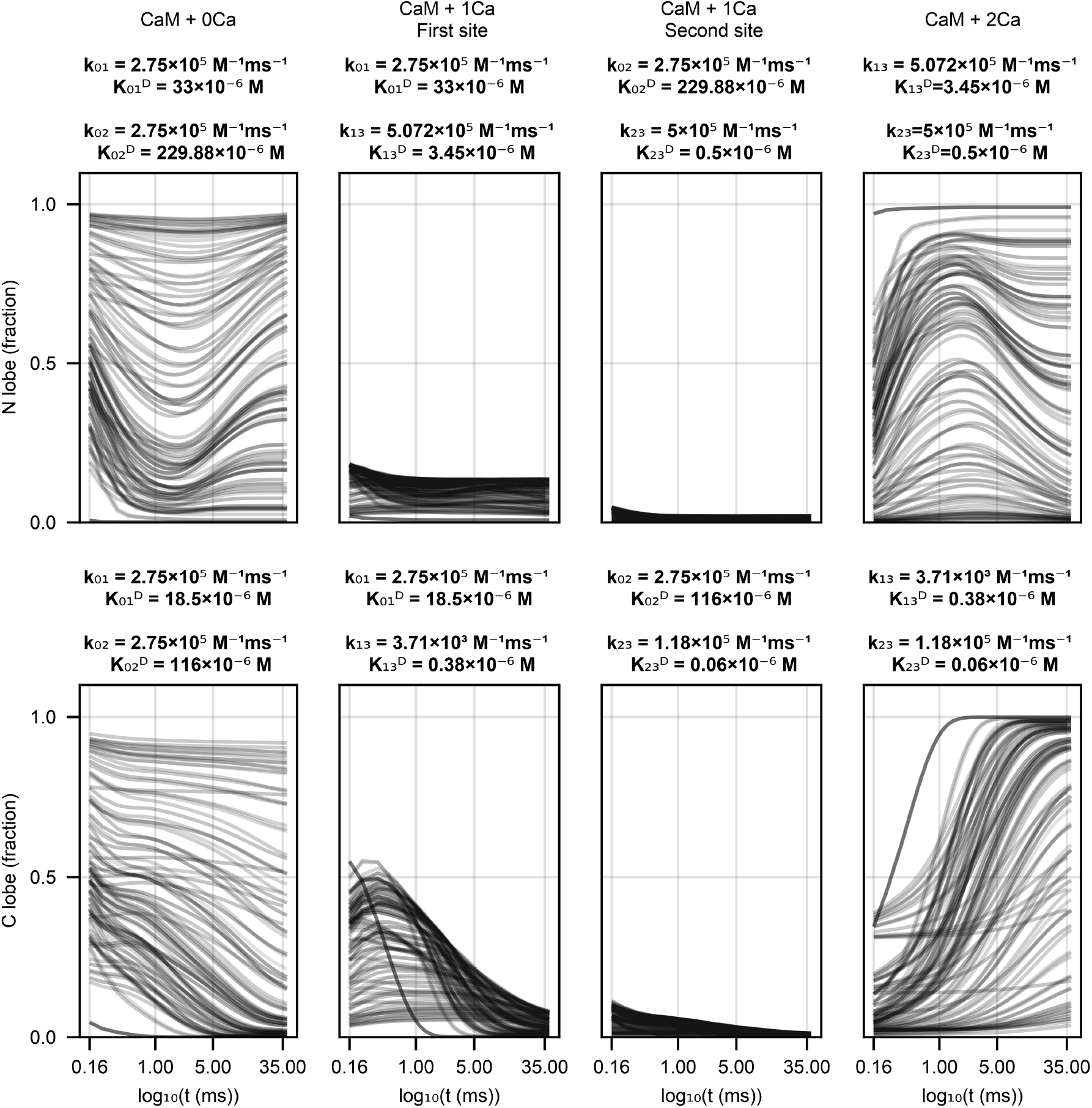
Scheme 6 behaviour with parameters from Byrne et al. [13] on dynamical data from Faas et al. [5]. Each line is one of the initial conditions (solution + uncaging strenght) Faas et al. used. Calmodulin has been normalized to total calmodulin used in an experiment, the first row shows N lobe dynamics, the second row shows C lobe dynamics. Each column shows a different calmodulin state – completely unbound (first column), Ca^2+^ bound to the first site on a lobe (second column), Ca^2+^ bound to the second site on a lobe (third column), Ca^2+^ bound to both sites of a lobe (fourth column). Above the plots we provide reaction rates from Byrne et al. [13] for the reactions a specific calmodulin state participates in. Note that time on the *x* axis is in log10 space to better show the initial dynamics.

Therefore, even though it is a theoretically appealing tool to reduce the number of calmodulin states, the quasi-steady state approximation is too inaccurate for the C lobe and results in significant errors in either calmodulin dynamics or equilibrium behaviour.

## Discussion

We used a rich dynamical [5] and equilibrium [11] data set to fit six calcium-calmodulin kinetic schemes from scratch in order to compare to published models. Our comparison resulted in a number of conclusions. First of all, the parameters we found, as opposed to the published ones, resulted in significantly better fits on our dataset (Table 1. Secondly, we showed that fully event-based schemes that do not utilize any features of the calmodulin physical structure (existence of C and N lobes) result in significantly worse generalization performance as measured via AIC (Fig 8). Thirdly, we investigated calmodulin signal integration properties by comparing our parameter fits to published reaction rates for different calcium-calmodulin schemes. Some schemes showed no Ca^2+^ signal integration in response to a stimulation protocol mimicking an empirically effective plasticity induction protocol highlighting the importance using more detailed calmodulin schemes (Fig 9). Fourthly, we calculated the partial correlations between our parameter fits (Fig 10). Partial correlations revealed that even with our data set, that is richer than anything used before, some parameters were correlated and therefore under-determined. Finally, we investigated the validity of the quasi-steady state approximation used in [12] and by using Faas et al. [5] data we showed that it is not accurate for the C lobe. We next discuss each of these conclusions individually.

First of all, model performance depends on the data which was used to parameterise it. Even though usage of multiple data sources to fit a calmodulin model is not new and was done in Pepke et al. [12], we are the first to combine a data source on calmodulin dynamics [5] and a data source on calmodulin equilibrium behaviour [11]. We used this combined data set to fit six different calcium-calmodulin kinetic schemes previously used in the literature. We then compared our parameters to the published ones which revealed that a significant number of calcium-calmodulin models used in the literature are parameterized sub-optimally (see Table 1). Most published models (except Faas et al. [5]) have relied on either only equilibrium data [11, 12, 40] or dynamical data obtained under significant methodological limitations – such as dead time in stopped flow fluorimetry or presence of other biochemical species [9, 13]. Undoubtedly, it would be unfair to criticize past work for operating under the limitations of the day, but that does not prevent models from becoming outdated (much as this work will be one day). Therefore, an important contribution in this paper are the improved model parameters for calcium-calmodulin models – the best performing parameter sets for each scheme are given in Table 3 (see S6 Appendix for all 20 parameter sets for each scheme).

**Table 3.**
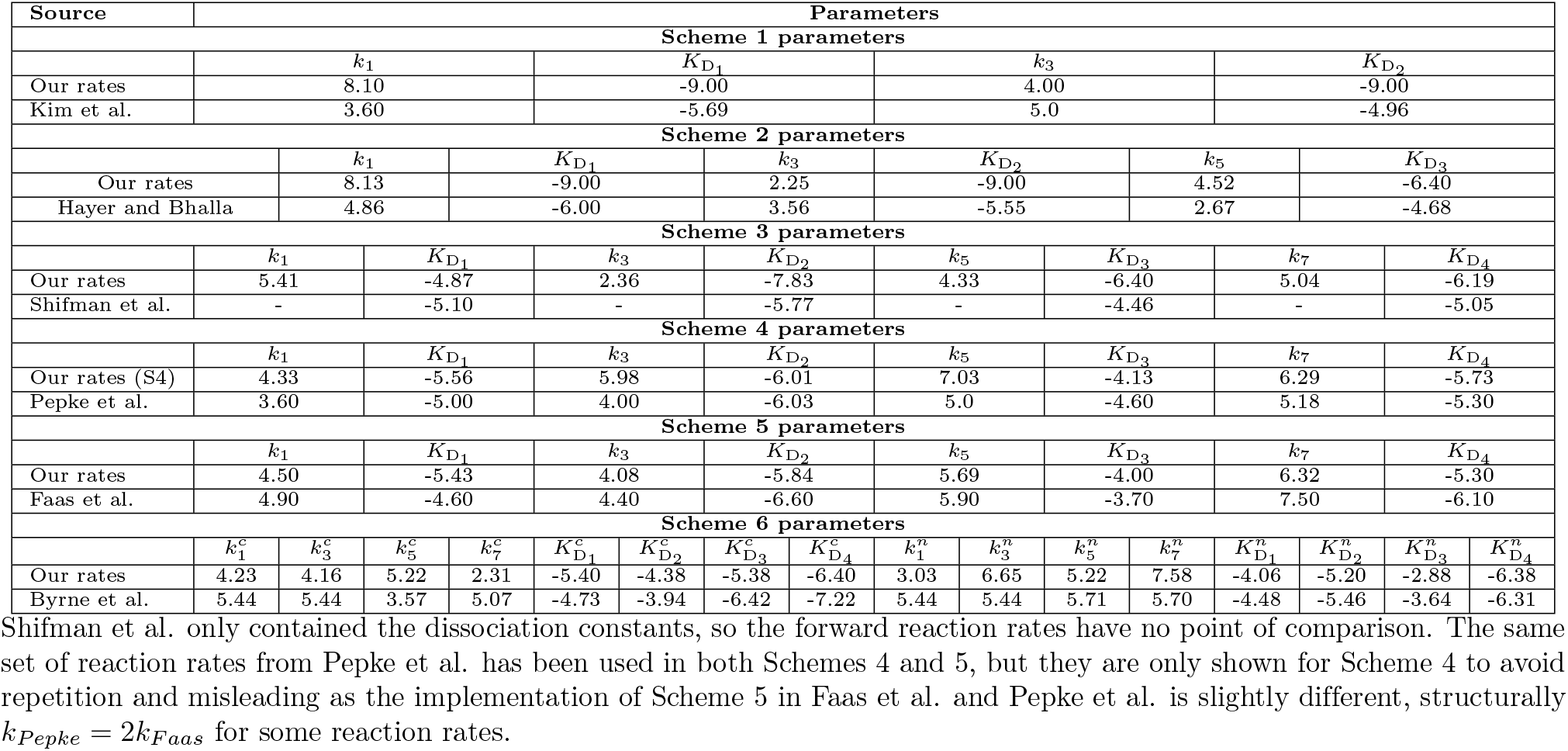
Our reaction rate sets that performed best on the test data and the published reaction rates from literature. All parameters are in log_10_, but are in different units, depending on the context: for second order reactions the forward rates are in M^-1^ms^-1^, dissociation constants in M, for third order reactions the forward rates are in M^-2^ms^-1^, dissociation constants in M^2^.

Secondly, our calmodulin model comparison uncovered discrepancies in performance between different kinetic schemes. The complexity of calmodulin schemes we investigated ranged from a model with three states and four parameters [9] to a model with eight states and sixteen parameters [13]. There were only two schemes (5 and 6, consisting of eight and sixteen parameters respectively) that were able to fit both sources of data well – both schemes modelled calmodulin lobes separately and consisted of individual, rather than lumped, Ca^2+^ binding reactions. Two further schemes (3 and 4), one of which modelled calmodulin lobes but not individual binding, another which modelled individual binding but not lobes, were able to fit dynamical data, but not equilibrium data, reasonably well. Both of these schemes consisted of eight parameters, same as one of the schemes that fit both sources of data well, indicating that the number of parameters is not the only factor necessary for an accurate calcium-calmodulin model. Finally, two of the simplest schemes (Schemes 1 and 2) that did not model calmodulin lobes and modelled Ca^2+^ binding as lumped reactions were not able to fit either the dynamical data or the equilibrium data well. These results, along with median AIC values (Table 2) lead to the second contribution of this paper – Scheme 6 is the most accurate calcium-calmoduling binding scheme and, compared to some simpler schemes, by a significant margin.

Thirdly, our results provide implications for models that include calmodulin. We investigated the Ca^2+^ integration properties of calmodulin in response to a realistic Ca^2+^ spike trains (see Fig 9). The biggest practical difference between our reaction rates and published ones is that there is a much more significant contribution from partially bound calmodulin species, rather than fully bound calmodulin. As shown in Shifman et al. [11], CaMKII can be activated by partially bound calmodulin. Moreover, calmodulin has many binding partners, such as Calcineurin [41], Phosphodiesterase

1 [42], Adenylyl cyclases 1 and 8 [43], Neurogranin [44, 45] and others [2]. Our results bring into question the accuracy of the results of publications where poorer performing schemes or parameterisations are used in larger models [9, 18, 19, 21–23, 46–49]. There are many ways to compensate for the poor performance of calmodulin scheme or parameters. For example, it is possible that in some cases the lack of calmodulin sensitivity to Ca^2+^ has been compensated for by an increased Ca^2+^ influx. However, for example Scheme 1 is used in [23] in a dynamical setting, stimulating their large model with many protein species with e. g. 180s of 5Hz or 1sec of 100Hz Ca^2+^ pulses. As our results show, the calmodulin Ca^2+^ integration properties are significantly different in this range when our reaction rates are used. Our third contribution is support to the hypothesis that partially bound calmodulin molecules arising in response to different Ca^2+^ stimuli is an additional dimension of signal encoding and propagation towards downstream pathways compared to spatial/concentration based fully bound calmodulin signalling. Future investigations into other calmodulin binding partners and their activation by partially bound calmodulin species would be able to falsify this hypothesis.

Fourthly, our results on the partial correlations between reaction rate constants form our fourth contribution – the call for more empirical investigations to test the distinctness of Ca^2+^ binding sites within a calmodulin lobe. Generally with increasing model complexity there were fewer correlations (except for Scheme 4, which had more than Scheme 3) between parameters, indicating the parameters were better determined by data. However, even for the most complex Scheme 6, there were some correlations between parameters within the same lobe. These correlations could only be eliminated by additional information on the properties of individual binding sites. Existing studies with mutations of individual calmodulin binding sites only include equilibrium measurements [11, 50, 51]. Since equilibrium behaviour only informs the ratio between the Ca^2+^ binding and unbinding rates, they are of limited usefulness in fitting. The closest to the necessary measurements were done in Faas et al. [5] where dynamical measurements with one inactive calmodulin lobe (either C or N) were made.

Finally, we investigated the validity of the quasi-steady state approximation used in [12]. Both Scheme 4, in which partially bound calmodulin species are not modelled due to the quasi-steady state approximation, and Scheme 5, which models them, can model calmodulin dynamics to a similar accuracy. The main difference between the schemes is in equilibrium behaviour, where in Scheme 4 the modelling of dynamics impedes modelling of steady state behaviour. These results imply that the quasi-steady state approximation used in Pepke et al. [12] does not hold in the context of the Faas et al. [5] data, at least not without significant decrease in the accuracy of model behaviour. Ideally, an empirical measurement of the occupancy of individual calmodulin sites in a dynamical setting would be definitive in falsifying this approximation. Unfortunately, such data does not exist therefore we used Scheme 6 with Byrne et al. [13] parameters (since they fit the data reasonably well) and simulated the fractional occupancy of individual calmodulin sites under [5] experimental conditions (see 11). These results support our fifth contribution – that the quasi-steady state approximation is not valid and results in a significant loss of accuracy, especially for the C lobe.

Having discussed the contributions of this paper we now reflect on their wider implications and practical reality of computational modelling. Suboptimal schemes or parameterisations of calcium-calmodulin models used in large models are a difficult challenge. It is not necessarily the case that the conclusions drawn from large models are made invalid. In large models it is likely possible to correct for the model-data mismatch arising due to inaccurate calmodulin behaviour via the parameters of reactions involving downstream molecules. This, however, may result in a panoply of different mechanistic hypotheses if different publications correct for these inaccuracies arising due to poor calmodulin models in different ways. A more co-ordinated community effort with some agreed upon set of model tests (such as the FindSim platform suggested by [52]), akin to continuous integration in GitHub, may be necessary to resolve such issues in the future and build performant large models.

Limited computational resources and the difficulty of writing large models mean that in some cases it may not be feasible to use a more detailed calmodulin scheme because of an exponential explosion in the number of species to be modelled and the subsequent increase in the computational cost of simulations. Rule-based modelling [53] with its “don’t care, don’t write” approach (only having to specify the features of a species which impact a reaction) allows models containing exponentially large numbers of complexes to be written down but may still be too computationally costly. Modeling is a complex task that involves many behind the scenes choices about acceptable trade-offs. Our results provide the information about the trade-offs in model accuracy being made when choosing one calmodulin scheme (or parameter set) over another.

In the final two paragraphs we discuss the methodology we used, the available alternatives and limitations. We used non-linear mixed effects model fitting algorithms implemented in `Pumas.jl` to fit the reaction rate constants of the different calcium-calmodulin kinetic schemes. There are many published pipelines for fitting reaction rates of kinetic schemes. For example, Eriksson et al. [54] propose and use a pipeline based on approximate Bayesian computation Markov Chain Monte Carlo (ABC-MCMC, using R-vines). MCMC approaches are powerful tools which benefit from inherently providing uncertainty on model parameters, rather than having to run optimization on different random seeds as was done in this study. However, they are generally much more computationally expensive. Another popular option is the Data2Dynamics toolbox [55], which streamlines construction of models of chemical reaction networks and modeling of experiments while leveraging ODE solving capabilities of MATLAB, along with stochastic optimization. However, there are few modern software packages that deal with non-linear mixed effects models (which were required due to the nature of the dynamical data in Faas et al. [5]). Of these packages `Pumas.jl` is currently the most performant one [24]. This is in part because `Pumas.jl` is implemented in the Julia programming language which contains state of the art ODE solving capabilities, outperforming its competitors in terms of speed by orders of magnitude (see benchmarks.sciml.ai).

Even with a powerful computational pipeline, there are still many nuances, practical considerations and limitations. For example, the length of the time series to which parameters are being fit impacts the complexity of the loss surface – the more points, the more complex it is [56]. Therefore, we downsampled the initial part of the dynamical data from Faas et al. [5] (see S3 Fig). However, invariably, downsampling results in loss of signal, therefore more performant downsampling techniques or multiple shooting based approaches may have resulted in even better fits. Moreover, we simplified the Ca^2+^ uncaging model used in [5] to make parameter optimization more stable. Also, [5] used Pockels cell delay (PCD) as the independent variable to predict the fraction of uncaged Ca^2+^ whereas we omitted this variable as it did not perform as well in practice. More data on the relationship between PCD and Ca^2+^ uncaging fraction would have allowed us to derive a better Ca^2+^ uncaging model that potentially could have improved model predictions with both published and our own reaction rates. Finally, in order to prevent training failures due to numerical instabilities in ODE solutions when using some schemes, we had to restrict the range of possible values taken by their reaction rate constants. Usage of novel ODE solvers capable of handling stiff systems is a potential avenue to remedy this limitation in future studies. Therefore, even with a more powerful software pipeline, some trial and error and practical trade-offs were necessary to fit our own parameters and efficiently and accurately compare different calmodulin models.

In conclusion, we believe that we have provided a number of important contributions that advance calcium-calmodulin modelling. We conducted a data-driven evaluation of both calcium-calmodulin kinetic schemes and parameter sets used in existing publications and showed which schemes or parameter sets performed poorly. It may be argued that behaviour of single molecules in large models matters less than the behaviour of the overall model. However, if large models are to be useful in predicting the behaviour of real biological systems, the individual molecules and their accurate generalization performance are of utmost importance.

## Supporting information

**S1 Appendix. Subset of data used and its splitting into training, validation and testing data sets**.

**S2 Table. Concentrations of species of different groups of solutions used in Faas et al. data**

**S3 Fig. Comparison of the original data set and the data with subsampled initial period**.

**S4 Appendix. Derivation of steady-state reaction rates for Scheme 4**.

**S5 Appendix. Published reaction rates used in this study**.

**S6 Appendix. Our parameter fits for all schemes**.

